# The shape of a defense-growth trade-off governs seasonal trait dynamics in natural phytoplankton

**DOI:** 10.1101/462622

**Authors:** Elias Ehrlich, Nadja J. Kath, Ursula Gaedke

**Author notes:** equally contributing authors (Elias Ehrlich, Nadja J. Kath).

## Abstract

Functional trait compositions of communities can adapt to altered environmental conditions ensuring community persistence. Theory predicts that the shape of trade-offs between traits crucially affects these trait dynamics, but its empirical verification from the field is missing. Here, we show how the shape of a defense-growth trade-off governs seasonal trait dynamics of a natural community, using high-frequency, long-term measurements of phytoplankton from Lake Constance. As expected from the lab-derived concave trade-off curve, we observed an alternating dominance of several fast-growing species with intermediate defense levels and gradual changes of the biomass-trait distribution due to seasonally changing grazing pressure. By combining data and modelling, we obtain mechanistic insights on the underlying fitness landscape, and show that low fitness differences can maintain trait variation along the trade-off curve. We provide firm evidence for a frequently assumed trade-off and conclude that quantifying its shape allows to understand environmentally driven trait changes within communities.

## Introduction

Identifying trade-offs between functional traits of species is central to ecology because it provides a fundamental basis to understand species coexistence and the trait composition of natural communities (Tilman 2000). Trade-offs emerge through physiological, energetic, behavioural or genetic constraints (Stearns 1989) and can be detected within one species (Barry 1994; Yoshida et al. 2004) as well as on the community level among different species sharing similar individual-level constraints (Tilman et al. 1982; Litchman et al. 2007). Such interspecific trade-offs promote species diversity and guide the way of community trait changes under altered environmental conditions (Kneitel and Chase 2004).

Theory indicates that it is the shape of the trade-off curve between two traits which determines species coexistence and how trait values change in response to environmental forcing (Levins 1962; Rueffler et al. 2004; Abrams 2006). We summarize the theory and specify predictions in Box 1. While theory revealing the importance of the shape of the trade-off curve for coexistence and trait dynamics is well developed (de Mazancourt and Dieckmann 2004; Jones et al. 2009; Ehrlich et al. 2017), its empirical verification has been left far behind. Two studies successfully tested the theory in small-scale lab experiments assembling different bacterial strains (Maharjan et al. 2013; Meyer et al. 2015). However, respective approaches from the field are lacking, leaving open the question how the shape of the trade-off curve affects the trait composition of natural communities. In this article, we combine theory and long-term field data, and show how the shape of a classical defense-growth trade-off affects seasonal trait dynamics of phytoplankton in a large European lake. Phytoplankton communities are well-suited for addressing this issue as important functional traits of phytoplankton have been measured in the lab revealing key trade-offs (Litchman and Klausmeier 2008; Pančić and Kiørboe 2018). Phytoplankton communities are extremely diverse spanning a large trait space (Weithoff 2003; Smith et al. 2005) indicating that trade-offs play a decisive role in maintaining trait variation. Furthermore, phytoplankton species have short generation times allowing for pronounced seasonal succession (Sommer et al. 2012). This offers the opportunity to observe species sorting in response to recurrently changing environmental conditions driving community trait dynamics.

Previous trait-based studies on phytoplankton communities already quantified trade-offs among different resource utilization traits (Litchman et al. 2007) and revealed how the trait composition of phytoplankton communities in different lakes and a marine system depended on light and nutrient conditions (Edwards et al. *2013a*,*b*). However, phytoplankton can be strongly top-down controlled selecting for phytoplankton defense, which was not considered in these studies but is likely to have a crucial effect on the seasonal trait dynamics (Sommer et al. 2012). Defense against predation often comes at the costs of a lower competiveness/ growth rate (Agrawal 1998; Pančić and Kiørboe 2018). This trade-off can mediate antagonistic effects of top down and bottom-up control on the trait composition. A large body of theory assumes such a trade-off (Abrams 1999; Tirok and Gaedke 2010; Klausmeier and Litchman 2012). However, there is no study which quantifies the shape of this trade-off and uses this information in combination with theoretical insights on trade-off curves (see Box 1 and Fig. 1) to explain how predation and abiotic conditions drive the trait dynamics and variation of natural communities.

**Figure 1.**
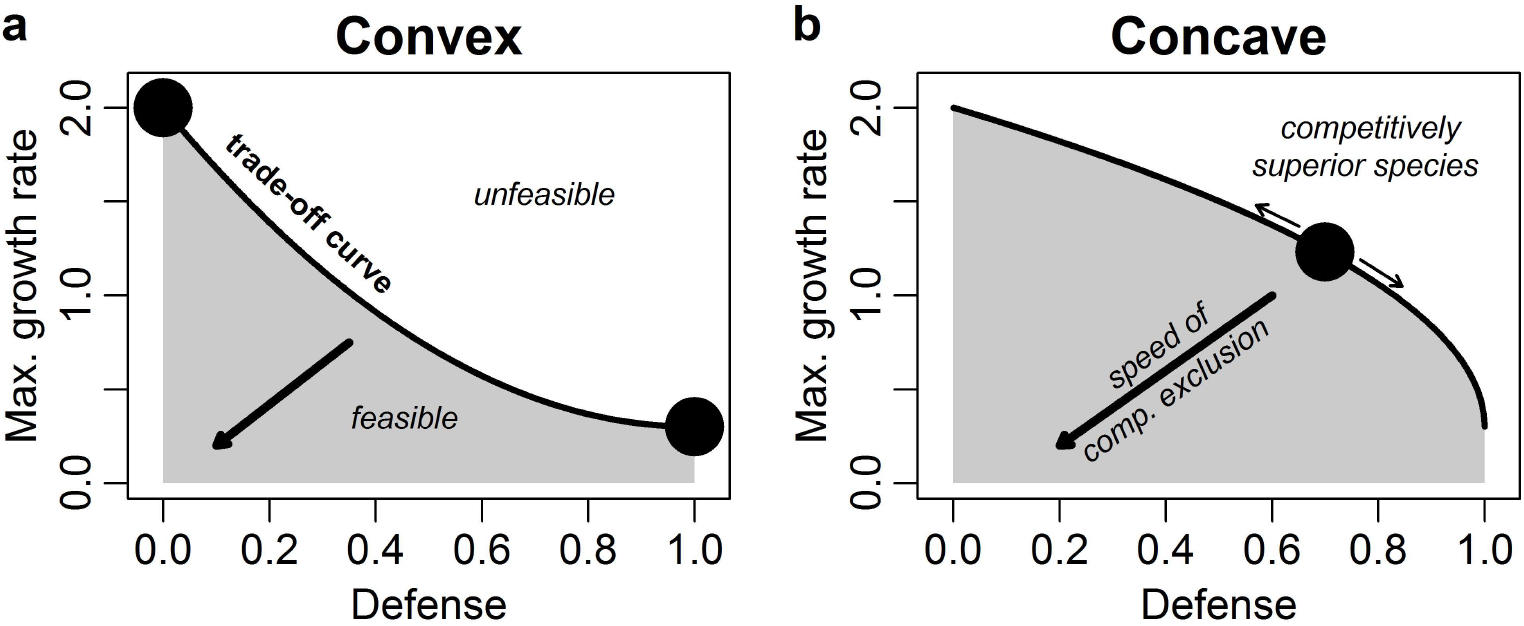
Competition outcome depending on the shape of the trade-off curve in a two dimensional trait space. The exemplary trade-off, shown here, is between defense against predation and maximum growth rate (*d^−^*^1^). The trade-off curve (black line) represents the boundary between the set of feasible (grey area) and unfeasible trait combinations (white area). (a) A convex trade-off curve allows for coexistence of two specialized species with extreme trait values (marked by circles) while the other feasible trait combinations are outcompeted in the long-term. (b) A concave trade-off curve promotes a species with intermediate trait values (circle) finally outcompeting the other species where it depends on the present environmental conditions which intermediate strategy is of maximal fitness as indicated by the thin arrows. The speed of competitive exclusion increases in the direction towards the unfavourable edge of the feasible trait space (low trait values) which is indicated by the thick arrows (a, b).

### Box 1 Theory on how the shape of a trade-off curve influences coexistence and trait dynamics

The trade-off curve is defined as the boundary of the set of feasible trait combinations, representing all possible phenotypes (Fig. 1, Rueffler et al. 2006). If the trade-off curve is convex, typically two specialized species coexist while all intermediate strategies are outcompeted in the long-term (given positive linear trait-fitness relationships, see Ehrlich et al. 2017) (Fig. 1a). If the trade-off curve is concave, only one species with intermediate trait values, determined by the present environmental conditions, is expected to survive in the long-term outcompeting all others (Fig. 1b). Nevertheless, on the short-term, more than one or two species may co-occur depending on the speed of competitive exclusion (Pedruski et al. 2015; Ehrlich et al. 2017) which is low for species with trait-combinations close to that of the species with maximal fitness and high for species at the unfavourable edge of the feasible trait space (Fig. 1a, b). Under directionally changing environmental conditions, the fitness maximum moves along a concave trade-off curve driving continued sorting of species with different trait values (Fig. 1b), e.g., an increasing grazing pressure promotes species with higher defense values at the cost of a decreasing maximum growth rate. In contrast, for convex trade-off curves, the fitness maxima usually stay at the extreme trait combinations (Fig. 1a) and only the biomass ratio between the specialized species is altered.

Here, we combine theory and field data to show the importance of the shape of a trade-off between defense and maximum growth rate for the trait dynamics of a natural, co-evolved phytoplankton community. We use a comprehensive data set of large, deep, mesotrophic Lake Constance where strong trophic interactions, vertical mixing and resource depletion are important alternating forcing factors of phytoplankton (Gaedke 1998). The data set comprises 21 years of taxonomically resolved, high-frequency measurements of phytoplankton and their grazers as well as measurements of abiotic factors (vertical mixing intensity and phosphorous concentration) which all undergo a highly repetitive seasonal succession. The considered major functional traits were defense against predation, maximum growth rate and phosphate affinity. We expect that each trait is promoted by certain environmental conditions: 1. Species with high defense levels are favored by high grazing pressure. 2. High maximum growth rates are generally advantageous as they imply high growth capabilities and can compensate for high losses due to deep vertical mixing which removes phytoplankton from the favorable, euphotic zone. 3. A high phosphate affinity is promoted under phosphorous depletion. The species trait values were taken from lab measurements mainly conducted with Lake Constance plankton (Bruggeman 2011) revealing a concave trade-off between defense and maximum growth rate, while there was no significant correlation of these traits with phosphate affinity. To obtain mechanistic insights on the underlying fitness landscape guiding trait changes, we developed a model parametrized with the found trade-off curve. We run numerical simulations to evaluate the favoured trait combinations and the speed of competitive exclusion of unfavorable trait combinations under different levels of grazing pressure, mimicking different seasonal field conditions. By linking theory, field data and modelling, we show that knowing the shape of the defense-growth trade-off is the key to understanding the ongoing trait changes and the maintenance of trait variation.

## Material and methods

### Study site and sampling

Upper Lake Constance (Bodensee) is a large (472 km^2^), deep (mean depth = 101 m), warm-monomictic, mesotrophic lake bordered by Germany, Switzerland and Austria. It has a well-mixed epilimnion and a large pelagic zone (Gaedke et al. 2002). Lake Constance underwent reoligotrophication during which the total phosphorous concentration declined 4-fold from 1979 to 1996 leading to an annual phytoplankton biomass and production decline by 50% and 25%, respectively (Gaedke 1998). This had no major effects on the biomass-trait distribution reported here and is thus not further considered.

Plankton sampling was conducted weekly during the growing season and approximately fortnightly in winter, culminating in a time series of 853 phytoplankton biomass measurements from 1979 to 1999 (for details see https://fred.igb-berlin.de/Lakebase). Phytoplankton counts and cell volume estimates were obtained using Utermöhl (1958) inverted microscopy and were converted into biomass based on a specific carbon to volume relationship (Menden-Deuer and Lessard 2000). Measurements were taken from the uppermost water layer between 0 and 20 m depth which roughly corresponds to the epilimnion and the euphotic zone. In this study, we considered the 36 most abundant morphotypes of phytoplankton (constituting 92% of total phytoplankton biomass) comprising individual species or higher taxonomic units that are functionally identical or very similar under the functional classification employed here. This guaranteed a consistent resolution of phytoplankton counts across years. Zooplankton was sampled with the same frequency as phytoplankton. Data for all major herbivorous zooplankton groups (ciliates, rotifers, cladocerans and copepods) were available from 1987 to 1996.

### Seasonal patterns

We subdivided the year into seven consecutive phases: late winter, early spring, late spring, clear-water phase (CWP), summer, autumn and early winter. Each phase was characterized by a well-defined combination of biotic and abiotic factors driving the phytoplankton community (Fig. 2): Strong vertical mixing implying high phytoplankton losses from the euphotic zone occurred during winter and partly early spring. Grazing pressure was most important during the CWP and summer, and declined towards autumn. Nutrient depletion was most relevant in summer and autumn when vertical mixing, supplying nutrients from larger depths, was absent.

**Figure 2.**
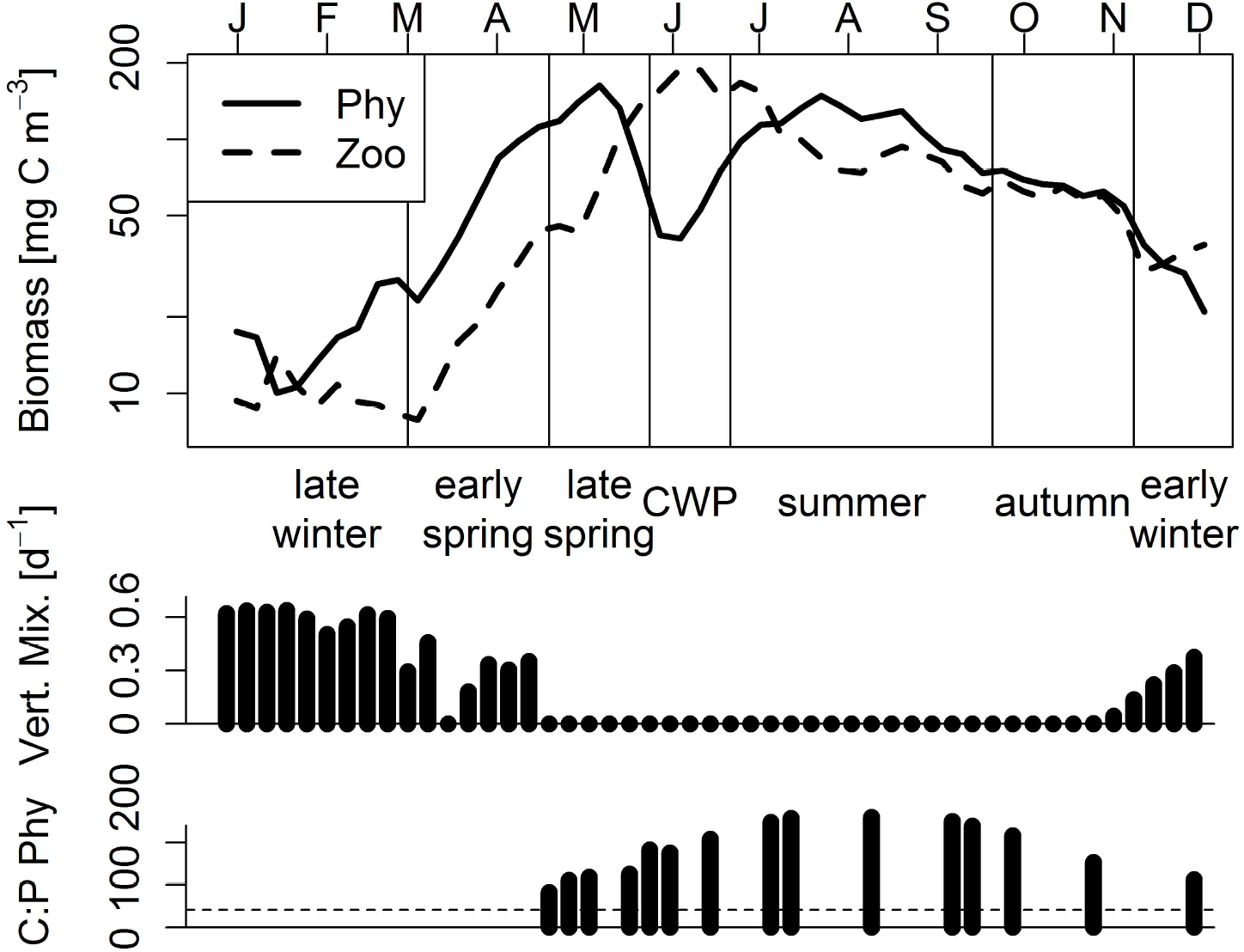
Inter-annual median biomass of phytoplankton (Phy) and herbivorous zooplankton (Zoo) in seven seasonal phases of a standardized year. The two bottom panels display the respective vertical mixing intensity which determines phytoplankton losses from the euphotic zone and the carbon to phosphorous (mass) ratio of phytoplankton indicating nutrient depletion (dashed line marks the Redfield ratio). For methodical details see Appendix 1.

### Trait data and trade-offs

The trait values for edibility, maximum growth rate and phosphate affinity of the 36 phytoplankton morphotypes were taken from Bruggeman (2011). Edibility was defined as the rate of prey consumption relative to the rate at which the favorite prey *Rhodomonas minuta* was consumed by *Daphnia* (Bruggeman 2011) which were very abundant prey and grazers in Lake Constance. We defined defense as 1 – edibility. All morphotypes, their assigned trait data and taxonomy are listed in Appendix 2. To detect potential pairwise trade-offs, we tested correlations between defense, maximum growth rate and phosphate affinity using the Spearman rank correlation.

### Model

We developed a simple model, based on Rosenzweig and MacArthur (1963), to show how the fitness landscape and the resulting biomass-trait distribution of a phytoplankton community differs under low (e.g., during early spring) and high grazing pressure (e.g., during summer). The model included N phytoplankton species which face a defense-growth trade-off, and one zooplankton group:

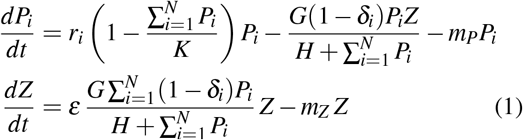

where *P_i_* represents the biomass of phytoplankton species *i*, *r_i_* the maximum growth rate, *δ_i_* the defense against zooplankton, *K* the carrying capacity and *m_P_* the natural mortality of phytoplankton. *Z* denotes the zooplankton biomass, *G* the maximum grazing rate, *H* the half-saturation constant, *ε* the conversion efficiency of phytoplankton biomass into zooplankton biomass and *m_z_* the mortality of zooplankton (for a detailed parameter description see Appendix 3). By changing *m_z_*, we can vary the importance of grazing pressure where a low *m_Z_* implies high zooplankton biomasses, that is, a high grazing pressure and vice versa. We assume a concave trade-off curve between *r_i_* and *δ_i_*, similar to the one found in the empirical data, and considered 184 different phytoplankton species with trait values spanning the whole feasible trait space. For details on the justification, parametrization, initialization and numerical integration of the model see Appendix 3.

## Results

The results section is divided into four parts: At first, we present insights on relevant trade-offs obtained from trait data for the morphotypes encountered in Lake Constance. Secondly, we describe the mean annual biomass-trait distribution of the phytoplankton community. Thirdly, we show how the biomass-trait distribution changes seasonally in response to altered environmental conditions. Finally, we compare the observed patterns with the model predictions.

### Trade-offs

The 36 dominant morphotypes co-occurring in large, deep Lake Constance covered a large range of values in defense *δ* and maximum growth rate *r* (Fig. 3). In general, a low *δ* was accompanied by a high *r* and vice versa, leading to a significant, negative correlation between them (*ρ* = *−*0.61, *p* = 10*−*5). The combination of high *δ* and high *r* was not found, despite its potentially high fitness, suggesting a physiological or energetic constraint. Nevertheless, many species with intermediate *δ* and high *r* (and also high *δ* and intermediate *r*) occurred implying a concave trade-off curve (Fig. 3). Trait combinations of low to intermediate *δ* and small *r* being of low fitness were not found indicating past competitive exclusion. We tested also for correlations of *δ* or *r* with phosphate affinity which were not significant. In the following, we focus on the trait dynamics in *δ* and *r* but add information on phosphate affinity when useful.

**Figure 3.**
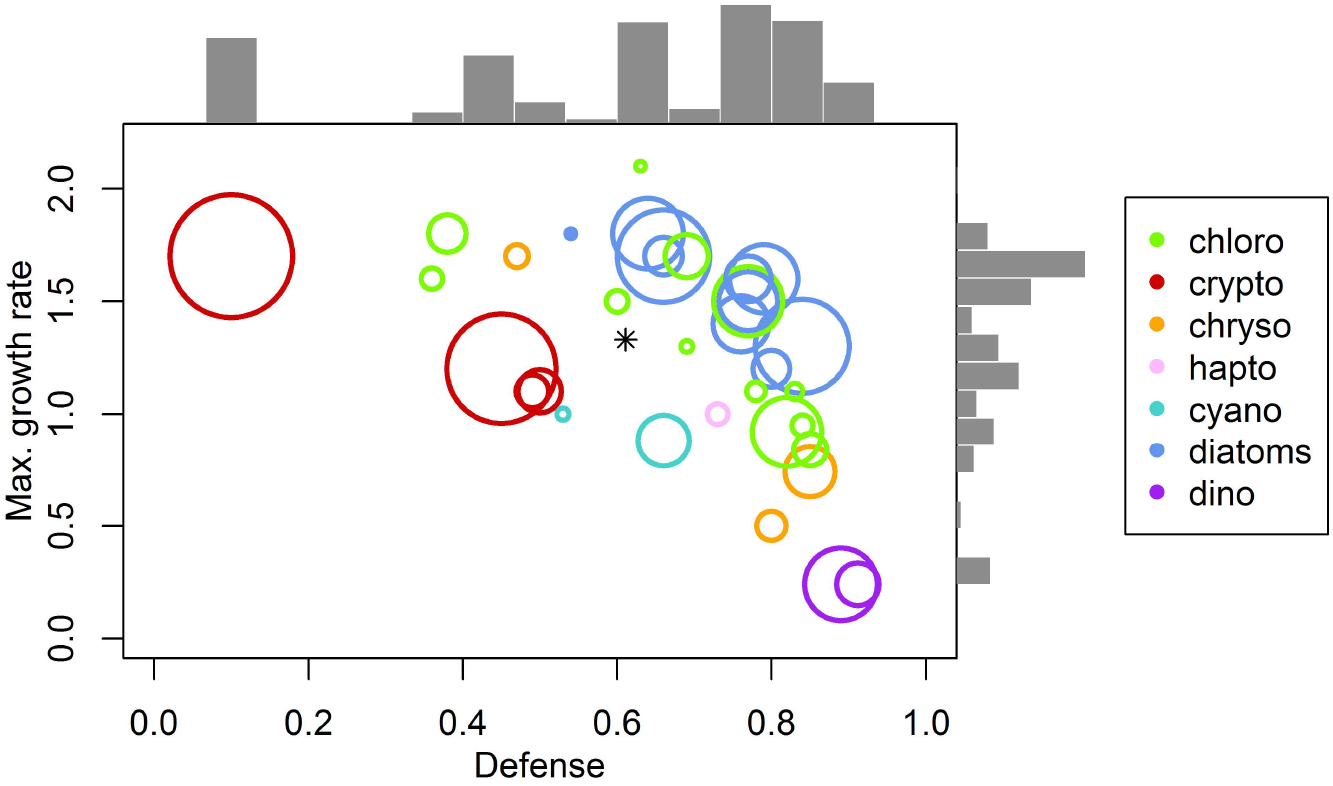
Defense *δ* and maximum growth rate *r* (*d^−^*^1^) of the 36 most abundant phytoplankton morphotypes in Lake Constance. Colors indicate different taxonomic groups, i.e., chlorophyta, cryptomonads, chrysophytes, haptophytes, cyanobacteria, diatoms and dinophytes. The area of the circles is scaled by the mean annual relative biomass of the morphotypes. The asterisk marks the biomass-weighted community mean trait values (*δ*̄, *r*̄). The bars display the relative biomass distribution along the two trait axes.

### Annual biomass-trait distribution

The mean annual biomass distribution within the trait space was obtained by weighting the morphotypes with their relative contribution to the total annual phytoplankton biomass (Fig. 3). The biomass was rather evenly distributed over the whole range of both traits with a maximum at intermediate to high values of *δ* and high *r*, caused by a cluster of different species of diatoms and chlorophytes. Considerable biomass occurs also at the extreme ends of the trait ranges (Fig. 3): *Rhodomonas* spp. with the lowest defense level and a high *r* constituted the highest annual share of biomass of an individual morphotype and occurred in almost every sample, and the highly defended but very slowly growing dinophytes exhibited intermediate mean annual relative biomasses. In general, for a given value of *δ*, the morphotypes with a higher *r* (i.e., higher fitness) dominated over those with lower *r* (Fig. 3). An exception to that was *Cryptomonas* spp. which only had an intermediate *r* despite its relatively low defence (*δ* =0.45) but the second highest mean annual relative biomass of an individual morphotype (Fig. 3). Remarkably, cryptomonads, chrysophytes, haplophytes and cyanobacteria generally had lower maximum growth rates compared to diatoms and chlorophytes given a certain defense level. For potential compensating advantages of these groups arising from further trait dimensions, see Discussion.

### Seasonal trait dynamics

Independent of the season, a large body of biomass lays along the concave trade-off and not below (Fig. 4). The biomass distribution within the trait space varied seasonally (Fig. 4) in line with pronounced changes of the major forcing factors of phytoplankton development (Fig. 2). These community trait changes can be tracked by considering the community mean trait values (*δ*̄, *r*̄) in the different seasonal phases, that is, the biomass-weighted average trait values among all morphotypes. In late winter and early spring, vertical mixing and the resulting high loss rate were the dominant driver of the phytoplankton community in deep Lake Constance while grazing pressure and nutrient depletion were very low (Fig. 2). Morphotypes with high *r* being able to compensate for high losses dominated, whereas morphotypes with low *r* and high *δ* were almost absent (Fig. 4a, b). This is reflected in the community trait means (late winter: *δ*̄ = 0.51, *r*̄ = 1.56 *d^−^*^1^; early spring: *δ*̄ = 0.52, *r*̄ = 1.57 *d^−^*^1^). During late spring, grazing pressure increased (Fig. 2) but did not initiate a shift of the overall biomass distribution towards higher *δ* (*δ*̄ = 0.48, *r*̄ = 1.55 *d^−^*^1^) (Fig. 4c). During the clear-water phase (CWP), the grazing pressure was at its annual maximum (Fig. 2). The mean community *r* decreased slightly (*r*̄ = 1.35 *d^−^*^1^) while the mean defense level did not change (*δ*̄ = 0.48) despite the high grazing pressure (Fig. 4d), probably due to a delayed numerical response of highly defended but slowly growing morphotypes. In summer, nutrient depletion and grazing pressure were the dominant drivers of phytoplankton. The biomass shifted towards morphotypes with intermediate or high *δ* and accordingly low *r* (Fig 4e, *δ*̄ = 0.69, *r*̄ = 1.18 *d^−^*^1^) but partially high phosphate affinity (Fig. 4e). In autumn, nutrient depletion and grazing were still mainly driving the phytoplankton community but declined compared to summer (Fig. 2). This resulted in a slight increase of morphotypes with lower *δ* and higher *r* (*δ*̄ = 0.62, *r*̄ = 1.28 *d^−^*^1^) (Fig. 4f). In early winter, vertical mixing again represented the most important driver. Morphotypes with high *r* and intermediate *δ* contributed a high share to the total phytoplankton biomass (Fig. 4g, *δ*̄ = 0.56, *r*̄ = 1.40 *d^−^*^1^).

**Figure 4.**
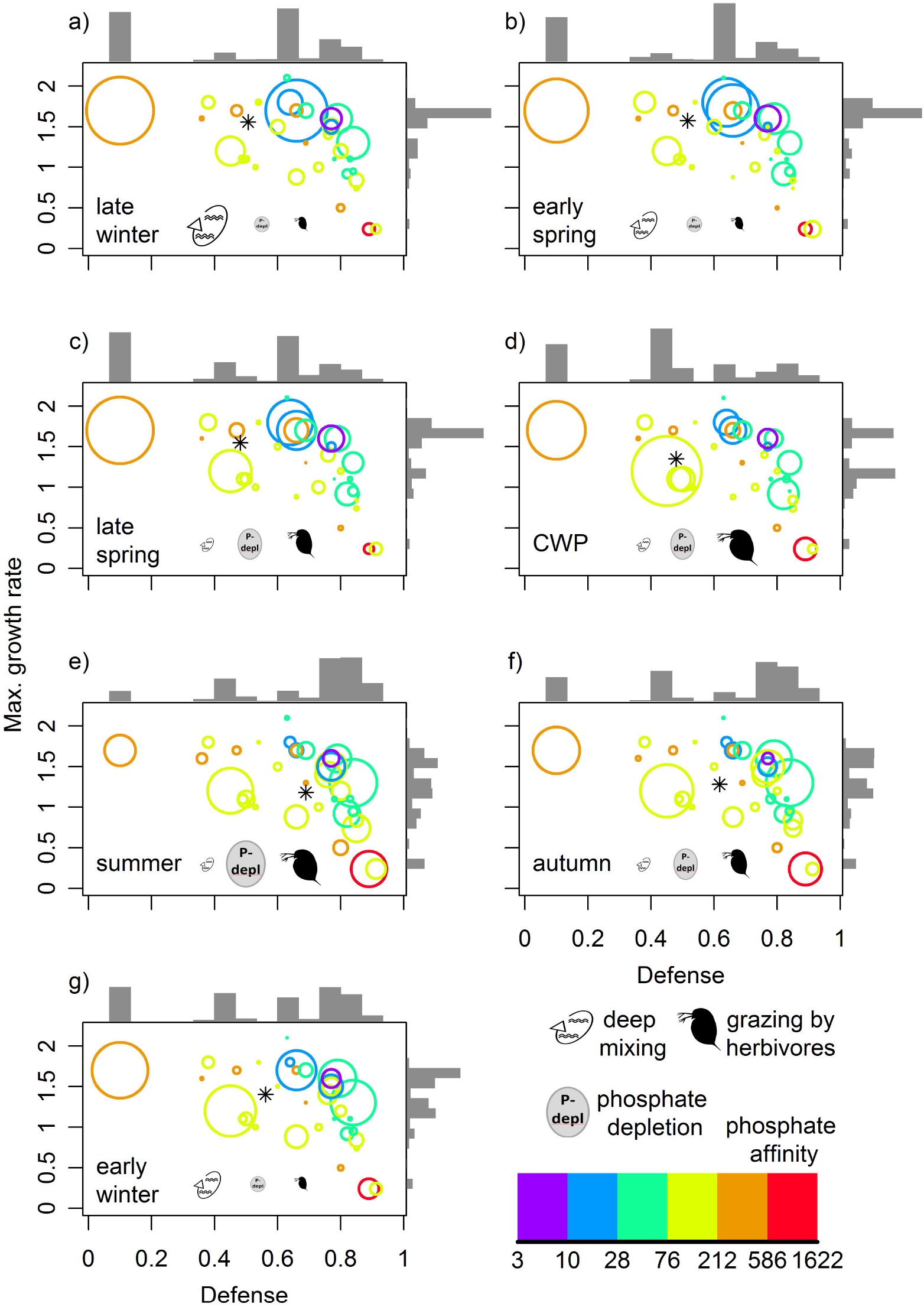
(a-g) Positions in the trait space of defense *δ* and maximum growth rate *r* (*d^−^*^1^) and mean relative bio-masses (scaling the area of the circles) of the 36 most abundant phytoplankton morphotypes in Lake Constance for the seven seasonal phases. The asterisks mark the respective biomass-weighted community mean trait values (*δ*̄, *r*̄). Colors indicate the phosphate affinity (*d^−^*^1^ *μmol^−^*^1^ *L*) of the individual morphotypes. The icons represent the dominant drivers of the phytoplankton community (vertical mixing, phosphate depletion, grazing by herbivores) and their size indicates their relative importance for phytoplankton net growth in each phase. The bars display the relative biomass distribution along the two trait axes in each phase.

The different morphotypes with intermediate or high *δ* dominating in different seasons had potentially different costs for their defense: In spring at deep vertical mixing and low nutrient depletion, the two dominant defended morphotypes had a high *r* but a very low phosphate affinity, while during summer with strong nutrient depletion and no deep vertical mixing, defended morphotypes had partly a more reduced *r* but a higher phosphate affinity (Fig. 4c, e). Hence, even if we found no significant correlation of *δ* or *r* with phosphate affinity within the whole community, some morphotypes may face a three-way trade-off. Nevertheless, the seasonal change in the community mean *δ* and *r* was stronger than for phosphate affinity (minimum during early spring: 215 *d^−^*^1^ *μmol^−^*^1^ *L*, maximum in autumn: 277 *d^−^*^1^ *μmol^−^*^1^ *L*) relative to the respective feasible trait range (3.6 *−* 1600 *d^−^*^1^ *μmol^−^*^1^ *L*).

### Model results

The model reproduced the general pattern in the empirical data, so that the favorable trait combinations shift from winter/spring (low grazing pressure) to summer/autumn (high grazing pressure) towards higher *δ_i_* at the cost of a lower *r_i_* (Fig. 4 a-f; 5a,b). For the given concave trade-off curve and set of trait combinations, the model predicted that two very similar species with intermediate *δ_i_* but high *r_i_* coexist in the long-term under low grazing pressure (Fig. 5a). Under high grazing pressure, the long-term outcome of the model was the survival of only one species with a high *δ_i_* but intermediate *r_i_* (Fig. 5a, b, for biomass dynamics see Appendix 4). When considering the short-term results of the model being more in line with the time scale relevant for the data of the different seasons, we found that many species along the concave trade-off curve (especially close to the fitness maximum) survived the first 50 to 100 days (Fig. 5a, b), in accordance with the observations (Fig. 4a-g). This holds in particular under low grazing pressure (Fig. 5a,b). Overall, the time until extinction was shorter under high grazing pressure due to the high mortality caused by abundant grazers (Fig. 5a,b). In general, the speed of competitive exclusion increased (fitness decreased) towards the unfavourable edge of the trait space (low *δ_i_*, low *r_i_*) where the slope of the fitness decrease depended on the degree of grazing pressure. Under high grazing pressure the fitness gradient was more perpendicular to the defense axis than under low grazing pressure (Fig. 5a,b).

**Figure 5.**
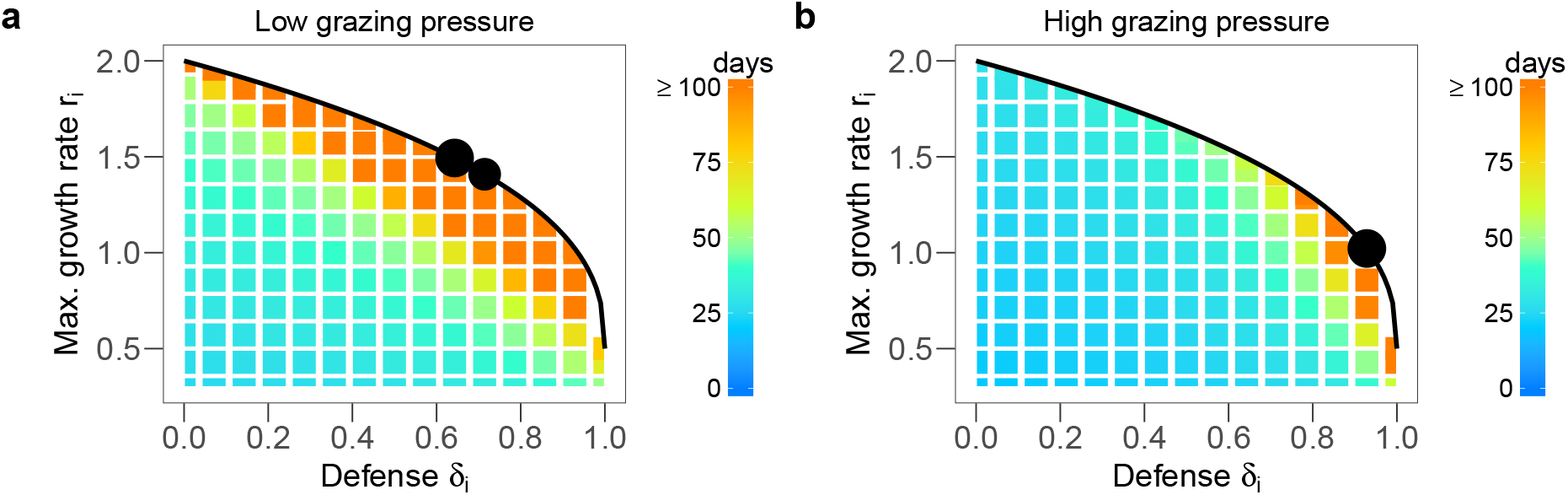
Model prediction on the competition outcome for a concave trade-off curve between defense *δ_i_* and maximum growth rate *r_i_* (black line) in the scenario of low grazing pressure on phytoplankton (*m_Z_* = 0.1 *d^−^*^1^) mimicking conditions in winter and spring (a), and the scenario of high grazing pressure (*m_Z_* = 0.02 *d^−^*^1^) during summer and autumn (b). The black dots denote the trait combinations of phytoplankton species which survive in the long-term, their size marks the mean relative biomass contribution between day 3000 and 4000 averaged among 100 simulations with randomized, different initial conditions (see Appendix 3). The colour grid displays the average time until extinction of the different trait combinations in the short-term, that is, within the first 100 days of the simulations. Note that trait combinations of species with *r_i_*-values below 0.3 *d^−^*^1^ (i.e., below the rate of natural mortality) are not displayed.

## Discussion

The 36 dominating phytoplankton morphotypes in Lake Constance faced a concave trade-off between defense and maximum growth rate. Theory predicts that concave trade-off curves promote species with intermediate strategies (see Box 1 and Fig. 1). Our data support this prediction as intermediately defended morphotypes with relatively high maximum growth rates constituted a high proportion of total annual phytoplankton biomass. Seasonal changes of the environmental conditions led to trait shifts in the phytoplankton community along the concave trade-off curve exactly as expected by the alternations in forcing factors. The model predicted a shift towards higher defense levels at the cost of lower maximum growth rates with increasing grazing pressure from spring to summer, as found in the data. Under constant conditions, the long-term outcome of the model was the survival of only one or two species with intermediate trait values. In contrast, we observed a high trait variation along the trade-off curve which can be explained by the short-term outcome of the model showing a slow competitive exclusion along the trade-off curve and shifting favorable trait combinations caused by seasonally changing conditions. The consideration of further trait dimensions such as phosphate affinity, which was not accounted for in the model, provide further explanation for the observed higher variation in defense and maximum growth rate. A high phosphate affinity, being relevant under nutrient depletion during summer, can compensate for low values of defense and maximum growth rate, and may allow the survival of the respective species.

The characterization of the trade-offs was based on trait data provided by Bruggeman (2011). He obtained trait values of the phytoplankton species from lab measurements and respective allometric calculations, mainly for the strains occurring in Lake Constance at the time of sampling. Uncertainties in trait values might arise from: 1.) Laboratory conditions which were not exactly the same as in the lake. 2.) Potential variation in the used allometric relationships (Taherzadeh et al. 2017). 3.) Phenotypic plasticity. However, all these uncertainties in the exact trait values were minor compared to the measured trait range. Hence, we argue that the lab-measured concave trade-off curve between defense and maximum growth rate adequately reflects the main trait composition of the phytoplankton community in Lake Constance.

Our model showed that, for a concave trade-off curve, two very similar species can stably coexist (Fig. 5a, Appendix 4). This is in contradiction with theory predicting the survival of only one species (see Box 1 and Fig. 1b). The two species have intermediate strategies close to the fitness maximum and coexist based on stabilizing mechanisms arising from their slight difference in defense and growth. However, this community is not evolutionary stable (Edwards et al. 2018). Given gradual evolution, we expect that one species would reach the exact fitness optimum via trait adaptation and out-compete the other for a concave trade-off curve (de Mazancourt and Dieckmann 2004; Rueffler et al. 2004). In nature, evolutionary processes do not always proceed gradually, i.e., with very small steps of trait adaptation, implying that reaching the exact fitness maximum can be unlikely. Hence, stable coexistence of similar but slightly different species close to the fitness maximum may be relevant in natural system.

In the model, only a low trait variation was maintained in the long-term based on a concave trade-off between defense and maximum growth rate. This is in contradiction with the empirical data showing a large trait variation including specialized species (highly defended or fast growing) and species having intermediate strategies. Abrams (2006) showed for a competition model that stable coexistence of two specialists using two different resources and one generalist is possible under asynchronous resource fluctuations. The species coexisted based on the relative non-linearity in their resource uptake functions. Such relative non-linearity enabling stable coexistence has not been found for the predator-prey model considered here. However, fitness-equalizing mechanisms may also promote species coexistence (Chesson 2000; Hubbell 2001). A fitness-equalizing trade-off implies that species with different trait values along the trade-off curve have equal fitness because the rate of loss determined by one trait (here: defense) and the rate of gain determined by the other trait (here: maximum growth rate) are exactly balancing (Ostling 2012). Trade-off curves are only fitness-equalizing for a specific shape equal to the shape of the fitness isoclines making fitness-equality improbable to occur in nature (Purves and Turnbull 2010). For the considered trade-off between defense and maximum growth rate, the fitness-equalizing case corresponds to a linear trade-off curve (Ehrlich et al. 2017, see colour gradient in Fig. 5 a,b) which was not found in the empirical data (see Fig. 3).

Even if the trade-off was not fitness-equalizing, we argue that low fitness differences along the concave trade-off curve allow for short-term coexistence due to slow competitive exclusion (see Fig. 5a, b). Given environmental fluctuations which continuously alter the selection regime, a low speed of competitive exclusion may promote coexistence even in the long-term (Huston 1979). In Lake Constance, the environmental conditions change seasonally and move the fitness maximum gradually along the trade-off curve from fast growing, intermediately defended species in early spring to slowly growing but more defended species in summer and then back in winter. Thus, several species along the trade-off curve have maximal fitness at different times of the year. This pattern of gradually moving fitness maxima is specific to concave trade-off curves and is not expected for convex ones (see Box 1, Fig. 1a and Appendix 5). We argue that long-term coexistence of numerous phytoplankton species along the concave trade-off curve is possible based on: 1.) their high production under respective favorable conditions, given their short generation times, 2.) their slow competitive exclusion under unfavorable conditions and 3.) their ability to form resting stages (Fryxell and of America 1983) causing a storage effect (Chesson 2000). Feasible but unfavorable trait combinations (i.e., low defense and low maximum growth rate) apart from the trade-off curve were quickly outcompeted in the model, providing an explanation for their absence in Lake Constance and in the whole data set of Bruggeman (2011)(see Appendix 2).

The maintenance of a high trait variation in the lake was likely also promoted by further niche dimensions and respective traits which were not considered in the model. For instance, a high phosphate affinity is beneficial under strong nutrient depletion during summer and autumn, and may explain why the defended dinophytes survive despite their very high defense costs regarding the maximum growth rate (Fig. 3). The undefended *Rhodomonas* ssp. also had a high phosphate affinity which sheds light on its observed high biomasses and very regular occurrence in spite of its maximum growth rate not exceeding the one of intermediately defended species. In addition to that, *Rhodomonas* spp. is able to use additional light spectra based on the red accessory photopigment phycoerythrin allowing photosynthesis at greater depths which is relevant year round due to vertical mixing and self-shading. The same holds for *Cryptomonas* spp. which also reached high biomasses irrespectively of its low maximum growth rate relative to its defense level. The cyanobacteria (*Anabaena* spp. and *Oscillatoria* spp.) also produce additional photo-pigments and further increase their fitness by buoyance regulation which probably compensate for their relative low maximum growth rates compared to diatoms with similar defense levels. Diatoms seem to have maximal fitness regarding their defense and maximum growth rate. However, they face disadvantages due to the production of shells implying an additional silica demand and causing high sedimentation rates which lead to lower net growth rates than expected from their maximum growth rate. This is less relevant under intensive vertical mixing. Mixotrophy represents an additional niche dimension promoting the highly defended, bacterivorous *Dinobryon* spp. which exhibits a very low maximum growth rate but is able to obtain relevant amounts of phosphate from bacterivory under phosphorous depletion in Lake Constance (Kamjunke et al. 2006). In general, phytoplankton species of the same taxonomic group cluster within the trait space (Fig. 3) indicating shared ecological strategies among closely related species.

Lake Constance has successfully served as a model system for large open water bodies including marine ones (Gaedke 1992). It exhibits a typical seasonal plankton succession driven by vertical mixing, grazing and nutrient limitation (Sommer et al. 2012). These environmental factors are also main drivers of marine phytoplankton which is ecologically similar to freshwater phytoplankton and may face similar trade-offs (Kilham and Hecky 1988). Trade-offs between defense and growth are also relevant in terrestrial plant communities, for example, grasslands (Lind et al. 2013). Even independent from the considered system with its environmental drivers and relevant trade-offs, our approach provides a general solution for obtaining mechanistic understanding of ongoing trait changes directly under field conditions.

Overall, to summarize, the identification of the major trade-off and its shape provided a remarkable key to understanding trait shifts and altering species composition in the phytoplankton community under seasonally changing environmental conditions. Although multiple trait dimensions might play a role, our results showed that defense and maximum growth rate represent key traits in Lake Constance where grazers are known to strongly impact phytoplankton net growth (Gaedke et al. 2002) and vertical mixing can cause high biomass losses from the euphotic zone (Gaedke et al. 1998). The maintenance of trait variation was linked to low fitness differences and the changing environment which continuously moved favorable trait combinations along the concave trade-off curve preventing competitive exclusion. Our study successfully explained major trait dynamics based on a model including only the two-dimensional, interspecific trade-off between defense and growth, and allowed to verify the theory on the shape of the trade-off curve in the field. Our findings revealed that the combination of trait and biomass data with simple models, involving the major trade-offs found in the data and information on their shape, represents a powerful approach to understanding trait dynamics and variation in natural communities.

## Acknowledgements

Many thanks to Alexander Wacker, Alice Boit and Guntram Weithoff for very constructive comments on an earlier version of the manuscript. We thank Michael Raatz for fruitful discussions on the results. This research was funded by the German Research Foundation (DFG, GA 401/26-1/2).

